# Functional Proteomics of ABC Importers Reveal Synchronization of Mechanisms, Cellular Abundances, and Counterintuitive Stoichiometries

**DOI:** 10.1101/2025.02.23.639702

**Authors:** Hiba Qasem Abdullah, Nurit Livnat Levanon, Michal Perach, Moti Grupper, Tamar Ziv, Oded Lewinson

## Abstract

Prokaryotes acquire essential nutrients primarily through ABC importers, consisting of an ATPase, a permease, and a substrate-binding protein. These importers are highly underrepresented in proteomic databases, limiting our knowledge about their cellular copy numbers, component stoichiometry, and the mechanistic implications of these parameters. We developed a tailored proteomic approach to compile the most comprehensive dataset to date of the E. coli ‘ABC importome’. Functional assays and analysis of deletion strains revealed novel mechanistic features, linking molecular mechanisms to cellular abundances, co-localization, and component stoichiometries. We observed 4-5 orders of magnitude variation in import system abundances, with copy numbers tuned to nutrient hierarchies essential for growth. Abundances of substrate-binding proteins are unrelated to their substrate binding affinities but are tightly, yet inversely, correlated with their interaction affinity with permeases. Counterintuitive component stoichiometries are crucial for function, offering insights into the design principles of multi-component protein systems, potentially extending beyond ABC importers.

**Teaser:** Bacteria’s nutrient absorption secrets unveiled: deciphering the complexity of ABC transporter systems required for optimal growth.

## Introduction

ABC transporters comprise one of the largest and oldest protein families of any proteome, with ∼120 members in higher plants, 49 in human, and ∼80 in prokaryotes. They harness the energy of ATP hydrolysis to move molecules across cell membranes (1,2). In humans, ABC transporters protect cells from toxic materials and deliver biomolecules to designated compartments (3,4). They are directly linked to human diseases and to tumor chemotherapy resistance (5,6). In prokaryotes, the majority of ABC transporters function as importers (hereafter ABC importers) where each ABC importer provides the main route for the high affinity acquisition of specific nutrient or a group of structurally similar nutrients. Under conditions of nutrient limitation, such as those encountered during host infection, their role is underscored. It is therefore not surprising that a large number of bacterial ABC importers have been identified as key virulence determinants (7,8).

The functional unit of an ABC transporter minimally consists of four domains: two transmembrane domains (TMDs) that form the translocation pathway and two cytoplasmic nucleotide-binding domains (NBDs) that provide energy via ATP hydrolysis. ABC transporters that function as importers additionally require a substrate binding protein (SBP) which binds the substrate and delivers it to its cognate transporter (9–11).

Knowledge of absolute protein levels (*i.e.,* copy number per cell and protein cellular concentrations) is essential for kinetic modeling of biological processes, for comparing concentrations of different proteins within or across samples or species and for quantifying protein-level adaptations to changes in internal or external conditions (12–16). As ABC transporters provide the main route for prokaryotic nutrient import any quantitative study of cellular metabolite influx or metabolism/anabolism balance must consider their copy number. For multi-component systems such as ABC importers, other parameters that need to be considered are component stoichiometry and component localization/co-localization (17–19).

Likely due to their membranal localization, ABC transporters are heavily underrepresented in current proteomic datasets (16,20–23). In addition, unlike other protein families (24–28), ABC transporters have never been specifically targeted by proteomic studies. Therefore, despite their clinical importance, we still lack a rigorous understanding of their copy number and stoichiometries and of the mechanistic implications of these parameters.

Herein, we developed a tailored protocol for the label-free tandem mass spectrometry (LC-MS^2^) quantification of ABC importers and compile a comprehensive dataset detailing their cellular copy numbers and component stoichiometries. Based on this proteomic data, we performed functional assays and identified novel mechanistic features of ABC importers, linking molecular mechanisms and physiological roles to cellular copy numbers and component stoichiometries.

## Results and discussion

### Compilation of the ‘ABC importome’ dataset

Of the 79 ABC transporters encoded by the *E. coli* genome, we focused on the 48 systems which were suggested to function as import systems (29). These 48 putative import systems are currently underrepresented in the 18 available *E. coli* proteomic datasets. Furthermore, reported abundances of numerous components exhibit variations of 100-1000-fold between datasets (21,30,31) rendering the integration of disparate datasets unfeasible. Consequently, our objective was to compile a unified dataset that offers a comprehensive overview of the relative abundances of distinct systems.

We used the *E. coli* K12 strain BW25113, the progenitor strain of the Keio clone collection (32), a strain utilized in multiple proteomic investigations (16,33–35). Cultures were cultivated to mid-exponential phase utilizing the common M9-glucose medium. Initially, LC-MS^2^ analysis samples were prepared following a standard protocol based on SDS/urea extraction and denaturation, followed by tryptic digestion (see methods for complete details).

From the list of 48 putative *E. coli* ABC import systems (29), we excluded from consideration two systems that have been demonstrated to function as exporters, and another which functions as a glucose-1-P isomerase (36–38), and see Table S1.

The remaining 45 systems, hereafter referred to as the ‘ABC importome’ (Table S1) consist of 157 proteins, comprising 47 SBPs, 48 NBDs, and 62 TMDs. This composition mirrors the monomeric nature of SBPs, the homomeric arrangement of the NBDs, and instances of heteromeric TMDs.

As observed by others (16,39), using the common urea-SDS-trypsin sample preparation protocol we failed to identify many of the transmembrane domains (Table S2). Missing transmembrane domains (TMDs) of special interest included those of the import systems for maltose (MalFGK), histidine (HisPQM), vitamin B_12_ (BtuCD) zinc (ZnuBC), and molybdate (MolBC). To increase the number of identified proteins we prepared membrane fractions from the same cell pellets. This enabled the additional identification of 14 components that were masked by the complexity of cell pellets sample. To further improve the identification, we tried a total of 59 protocol modifications which included proteolytic cleavage with chymotrypsin, pepsin, LysC, WalP, or Proanalse, using n-Dodecyl-beta-Maltoside (DDM) or n-Decyl-beta-Maltoside (DM) instead of SDS, and washing membrane fractions with high salt, EDTA, or sodium bicarbonate (see methods for full details).

Table S2 lists the components of the ‘ABC importome’ that were identified by several of these alternative protocols. Of the different sample preparations we tried, the most significant improvement was achieved by washing the membranes with 100 mM sodium bicarbonate or using chymotrypsin instead of trypsin (7 and 9 additional components identified, respectively).

We now wished to combine the datasets obtained using the different sample preparation methods. For this, we first analyzed the correlation between them. As shown, we observed excellent correlations between technical and biological replicates (Figure S1A and S1B, respectively). Furthermore, a strong correlation was evident between the intensities of membrane proteins measured in whole cells and those in membrane fractions prepared from the same cells (Figure S1C), as well as between membrane protein intensities measured in membrane fractions and those measured in the same membranes washed in 100 mM sodium bicarbonate (Figure S1D). In contrast, we observed low to moderate correlations between samples digested with trypsin and those digested with chymotrypsin, LysC, pepsin, or Proanalse (Figure S2A-D). Similarly, we noted low correlations between samples extracted with SDS and those extracted with n-Dodecyl-β-D-maltoside (DDM), n-Decyl-β-Maltoside (DM), or a mixture of DM and DDM (Figure S3A-C). Based on these findings, we concluded that MS spectra obtained from biological replicates, membrane fractions, and membrane fractions washed with sodium bicarbonate can be integrated to form a unified dataset. In contrast, spectra acquired using alternative proteases or detergents could not be reliably combined.

Subsequently, we determined the fractional mass for each protein by utilizing its relative iBAQ value (*riBAQ*,(40,41)) and the known total protein content in the sample. These fractional masses were then converted to copy numbers employing the molecular weights of the proteins and Avogadro’s number. The ABC importome components were integrated into a consolidated dataset using the slopes of the correlation curves (Figure S1A-D), and cellular copy numbers were derived by dividing the total copy number of each protein by the number of cells as determined through colony counting. The complete mathematical expressions for these conversions are provided in the methods section. We then compared our estimated cellular copy numbers, derived solely from label-free LC-MS2, with those obtained using internal standards. In a previous study (16), the Heinemann group determined the absolute quantities of 41 *E. coli* proteins using heavy labeled reference peptides. By combining this data with the iBAQ approach, they quantified the entire *E. coli* proteome across 22 different growth media, including the M9-Glucose medium used in our study. Notably, both studies used the same *E. coli* strain (BW25113). For this comparison, we focused on substrate binding proteins (SBPs) since, unlike the transmembrane domains (TMDs) and nucleotide-binding domains (NBDs), identifying SBPs did not require additional sample processing steps (*e.g.,* preparation of membrane fractions and membrane washing with sodium carbonate), which were only applied in our study and not by the Heinemann group. Figure S4 shows the correlation between the cellular copy numbers of SBPs identified in the two studies, indicating a high correlation. Nevertheless, since we did not use in our study labeled internal reference peptides, for the subsequent analysis, we consider only order-of-magnitude differences between samples and systems. Our final compiled dataset (Table S3) presents the cellular copy number of 42 SBPs, 38 TMDs, and 33 NBDs, encompassing ∼72% (113 out of 157) of the ‘ABC importome’ components. To the best of our knowledge, this is the most comprehensive coverage of ABC transporters published to date.

### Abundances and stoichiometries of the ‘ABC importome’

Figure 1 shows the cellular copy numbers of the substrate binding proteins (SBPs) we identified, and the first striking feature that we noticed was the large variation in their abundances, spanning over more than four orders of magnitude (from nM to low mM concentrations). The most abundant SBPs, with approximately 20,000 copies per cell (resulting in a periplasmic concentration of ∼0.1 mM), were LivJ (Leu/Ile/Val SBP) and MetQ (L-Met SBP). In contrast, the least abundant SBPs, with an average of 1-3 molecules per cell (equating to a periplasmic concentration of approximately 2-6 nM), were BtuF and FhuD (vitamin B12 and ferric hydroxamate SBPs, respectively). We hypothesized that this significant variability might be related to the substrate binding affinity, where high abundances could potentially compensate for low substrate-binding affinities. However, as illustrated in Figure 2A, we did not observe such a correlation. For instance, the highly abundant SBP, MetQ, exhibits a very high affinity for methionine (∼0.2 nM, (42)), whereas FliY, which is approximately four-fold less abundant, binds cysteine/cystine with a 50,000-fold lower affinity of ∼ 10 μM (43).

**Figure 1.**
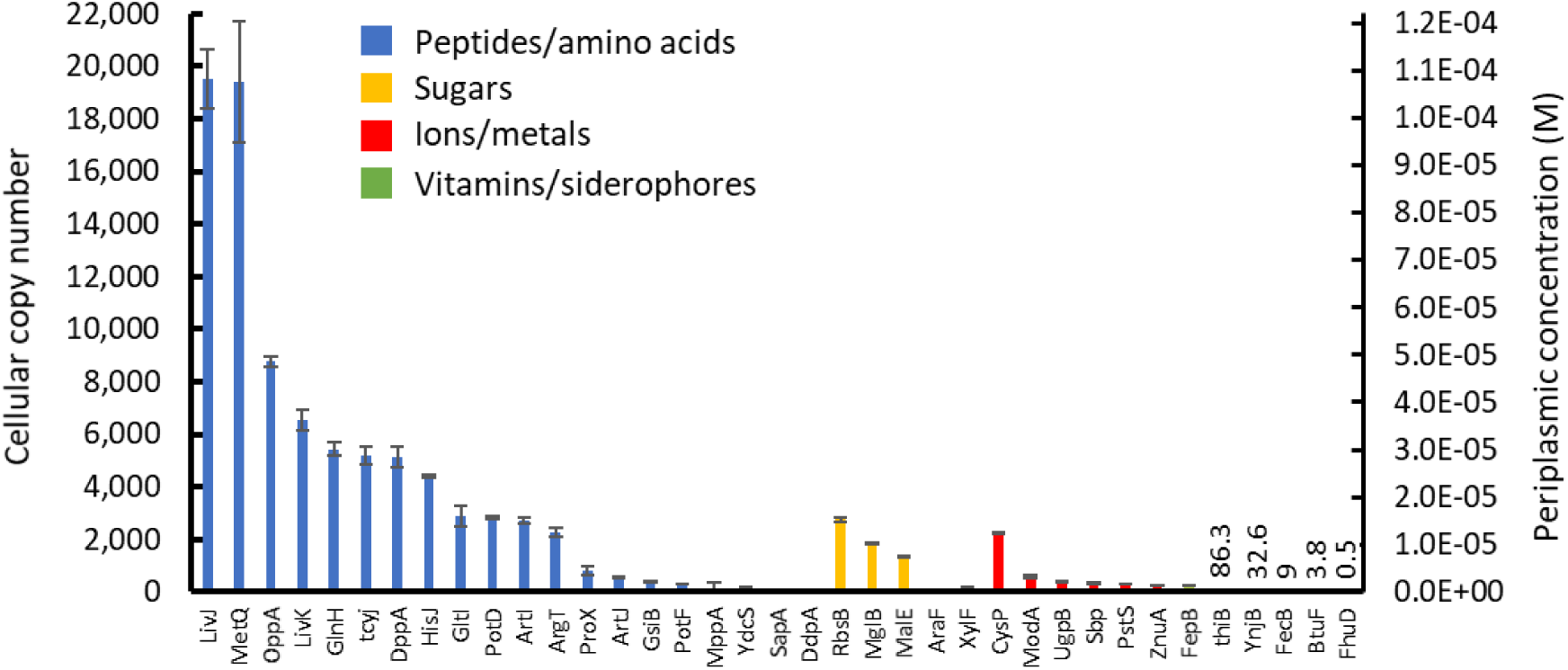
Segregation of the SBPs according to substrate class. Shown are the cellular copy numbers of 37 substrate binding proteins (SBPs), color coded as indicated according to their substrate group. Results are means of biological triplicates and error bars represent standard deviations.

**Figure 2:**
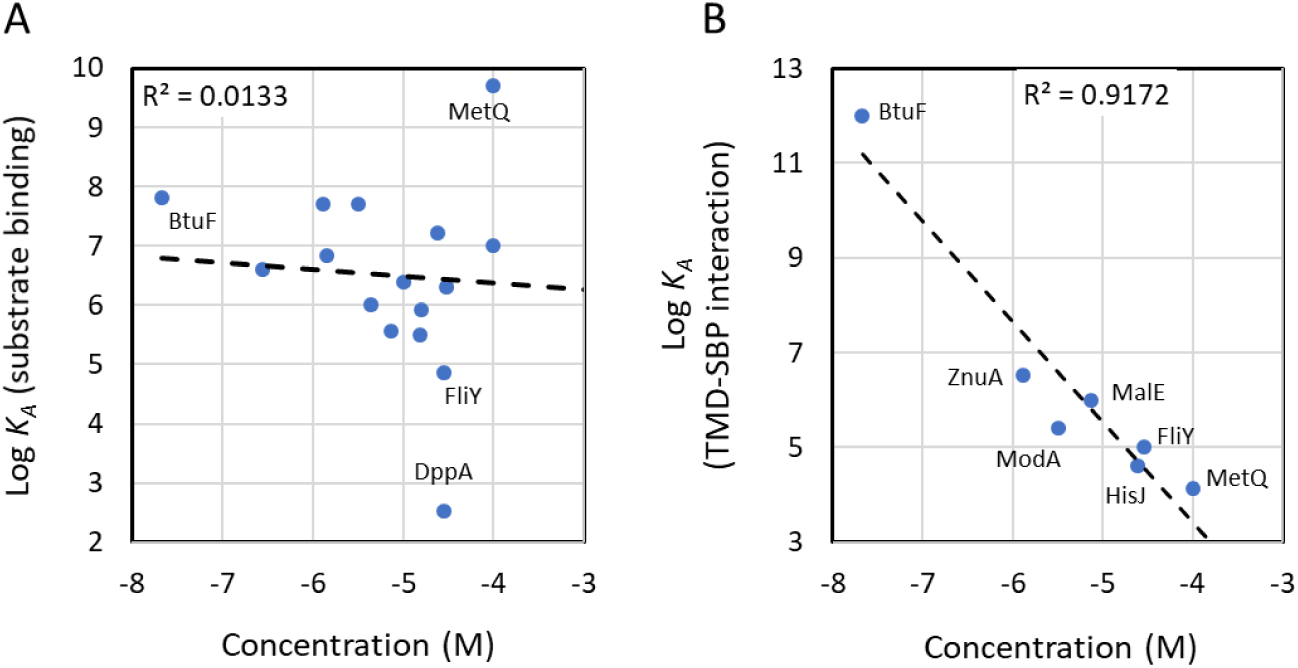
Correlation between SBP Abundances and binding Affinities. Shown are the correlations between the periplasmic concentrations of Substrate Binding Proteins (SBPs) and their substrate binding affinities (A) or their affinity towards cognate transporters (B). The dashed line represents the linear fit, and the goodness of fit is indicated.

An alternative driving force that may shape the abundance of the SBPs is their affinity of interaction with their cognate transporters, with SBPs that interact with their cognate transporters with low or high affinity present at high or low concentrations, respectively. Only seven values of SBP-transporter interaction affinity have been reported to date (44–50). However, despite this paucity of information we observed a clear correlation between the SBP-transporter interaction affinities and copy number, with high interaction affinities strongly correlating with SBPs of low abundance and low interaction affinities strongly correlating with high SBPs abundances (Figure 2B). It is difficult to resolve the chicken and the egg in this phenomenon: do low SBP abundances dictate high interaction affinities with the transporter or did high interaction affinities provide a selective pressure for reduced abundances. Whatever is the cause-effect relation, these two variables seem to be tightly linked.

In terms of abundance, the SBPs segregate into four distinct groups (Figure 1): The most abundant SBPs are those binding peptides and amino acids. This observation aligns with the high cellular demand for amino acids (51) and with findings indicating that protein synthesis is the energetically most demanding cellular process (52). The second most abundant group consists of SBPs binding sugars, which are also required in substantial amounts. As glucose served as the carbon source for growing the cultures, the lower abundance of sugar-binding SBPs may result from carbohydrate-mediated metabolic suppression (53). The third most abundant group comprises SBPs for ions (e.g., phosphate, sulfate, zinc), which are needed in smaller quantities (54). The least abundant SBPs are those for vitamins and siderophores, substances required in minute amounts. These results demonstrate that, in addition to transporter-SBP interaction affinity, substrate demand also influences the abundances of SBPs.

Similar to the SBPs, the abundances of the NBDs exhibited significant variation, ranging from approximately 2500 copies/cell (GlnQ, involved in glutamine uptake) to around 1 copy/cell (FecE, participating in Fe^3+^-dicitrate uptake). Their abundance hierarchy was comparable (though not identical) to that of the SBPs, i.e., peptides/amino acids > ions > sugars > vitamins/siderophores (Figure 3A). The TMDs displayed an identical abundance hierarchy, but their estimated cellular numbers were considerably lower than those of the NBDs (Figure 3B), which may be interpreted to suggest a stoichiometric excess of the latter. However, we could not detect any significant amount of the NBDs in the cytosolic fractions, where their abundance was similar to that of *bona fide* membrane proteins, suggesting that these remnants of “cytosolic” NBDs likely stem from incomplete removal of membrane fragments from the soluble fraction. As shown later (Figure 7) the NBDs are membrane associated in a TMD-dependent manner. We therefore suggest that that the apparent stoichiometric excess of the NBDs over the TMDs arises from the technical limitations of membrane protein quantification by LC-MS^2^ (55,56).

**Figure 3.**
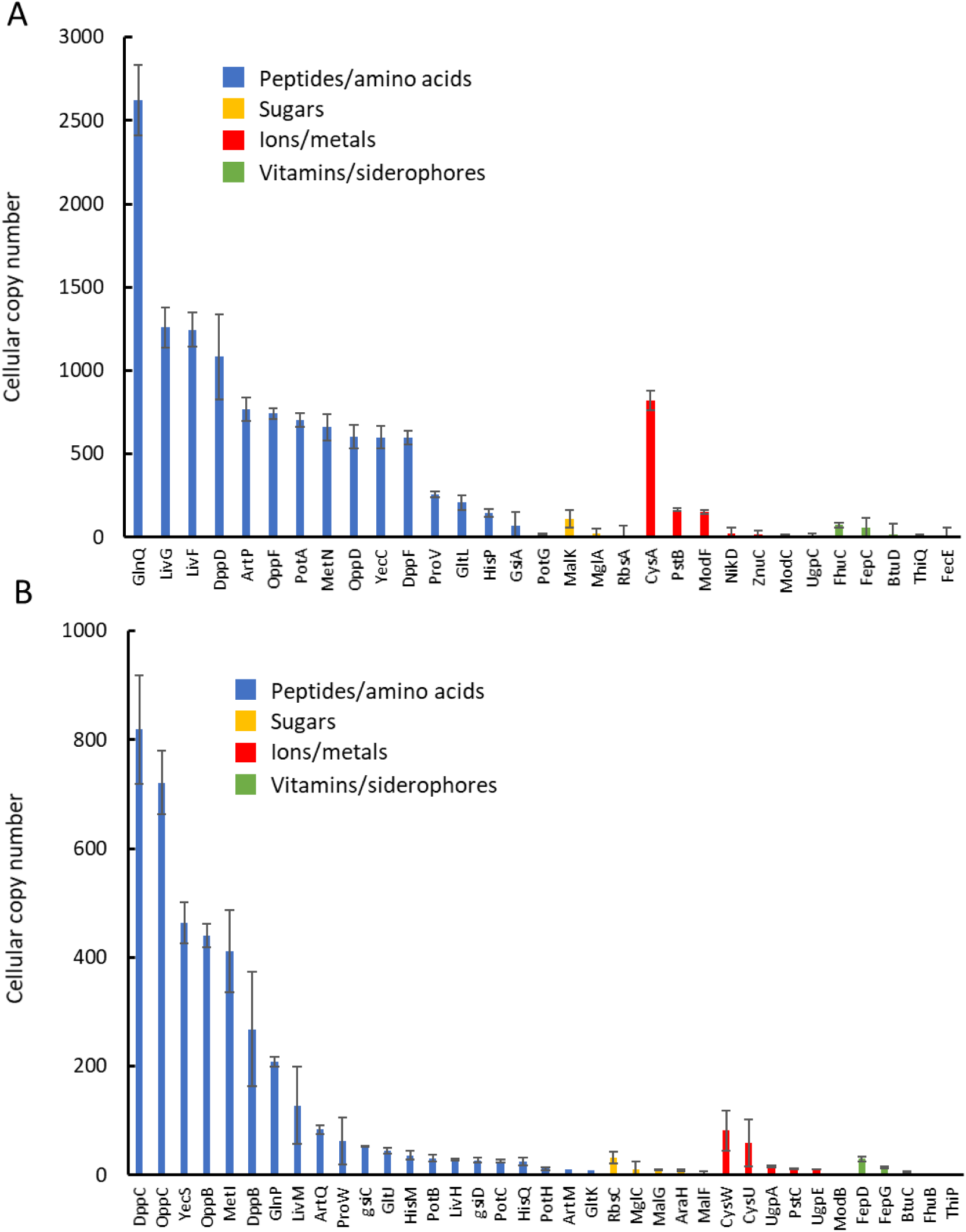
Cellular abundances of the Nucleotide Binding and Transmembrane Dominos. Shown are the cellular copy numbers of 30 NBDs (A) and 37 TMDs (B), color coded as indicated according to their substrate group. Results are means of biological triplicates and error bars represent standard deviations.

Another clear trend that emerges concerns the SBP-transporter stoichiometry: In all the systems belonging to the Type-I subgroup of ABC importers (10,57,58), i.e., systems that import peptides, amino acids, and sugars, we observed the SBP in stoichiometric excess relative to the transporter (Figure 4). Similar stoichiometric ratios have been deduced from measuring protein synthesis rates (59). In contrast, this ratio was reversed in all Type-II systems (systems that import siderophores and vitamins), and the transporter was found in stoichiometric excess relative to the SBP (Figure 4). This is a very surprising observation that raises questions about the relevance of in vitro transport assays conducted with such systems, which assume that the SBP-transporter stoichiometric ratio is >>1 (50,60–63). The functional implications of this surprising component stoichiometry in analyzed in the final results section of this report.

**Figure 4.**
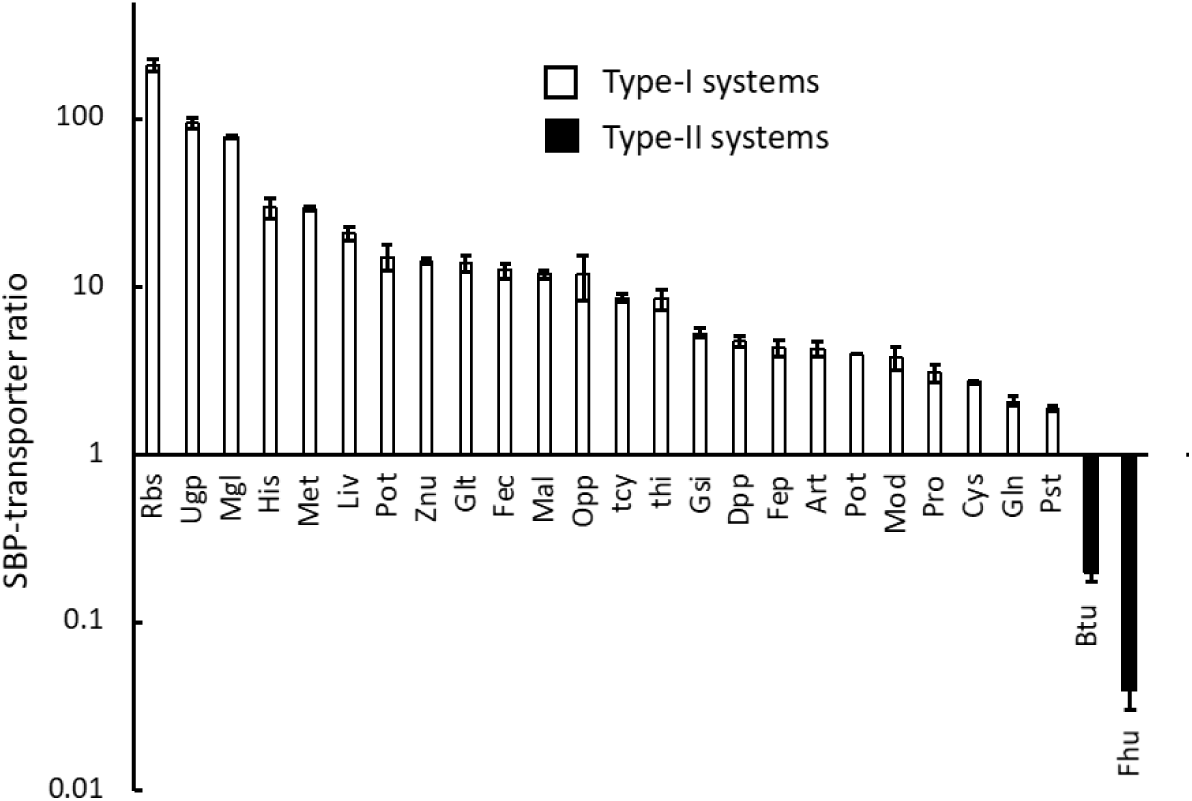
Different SBP: transporter stoichiometries in Type I and II systems. Shown are representative SBP-transporter stoichiometries determined for Type I (open bars) and II systems (full bars). Results are means of biological triplicates and error bars represent standard deviations.

When comparing whole-cell lysates to membrane fractions prepared from the same cells, we observed a 5-6-fold enrichment in the fractional abundances of the TMDs (Figure S1C). This enrichment factor is somewhat greater than the one expected based on the fraction of membrane proteins in *E. coli* (20-25%, (64)) and possibly reflects the non-monotonous technical improvement in detecting membrane proteins associated with reduced sample complexity (65). In contrast, the SBPs were heavily under-represented in the membrane fraction. This is expected since the periplasm is discarded during the membrane preparation. However, there were three exceptions to this trend, and the SBPs MetQ, BtuF, and FhuD were enriched in the membrane fraction (Figure 5). For MetQ, the SBP of the methionine import system, this is not surprising since MetQ is uncharacteristically tethered to the membrane via a lipid anchor (66,67). BtuF and FhuD on the other hand are “regular” periplasmic proteins that do not contain a membrane anchoring domain and presumably freely diffuse in the periplasm. This phenomenon is analyzed in detail in the next section.

**Figure 5.**
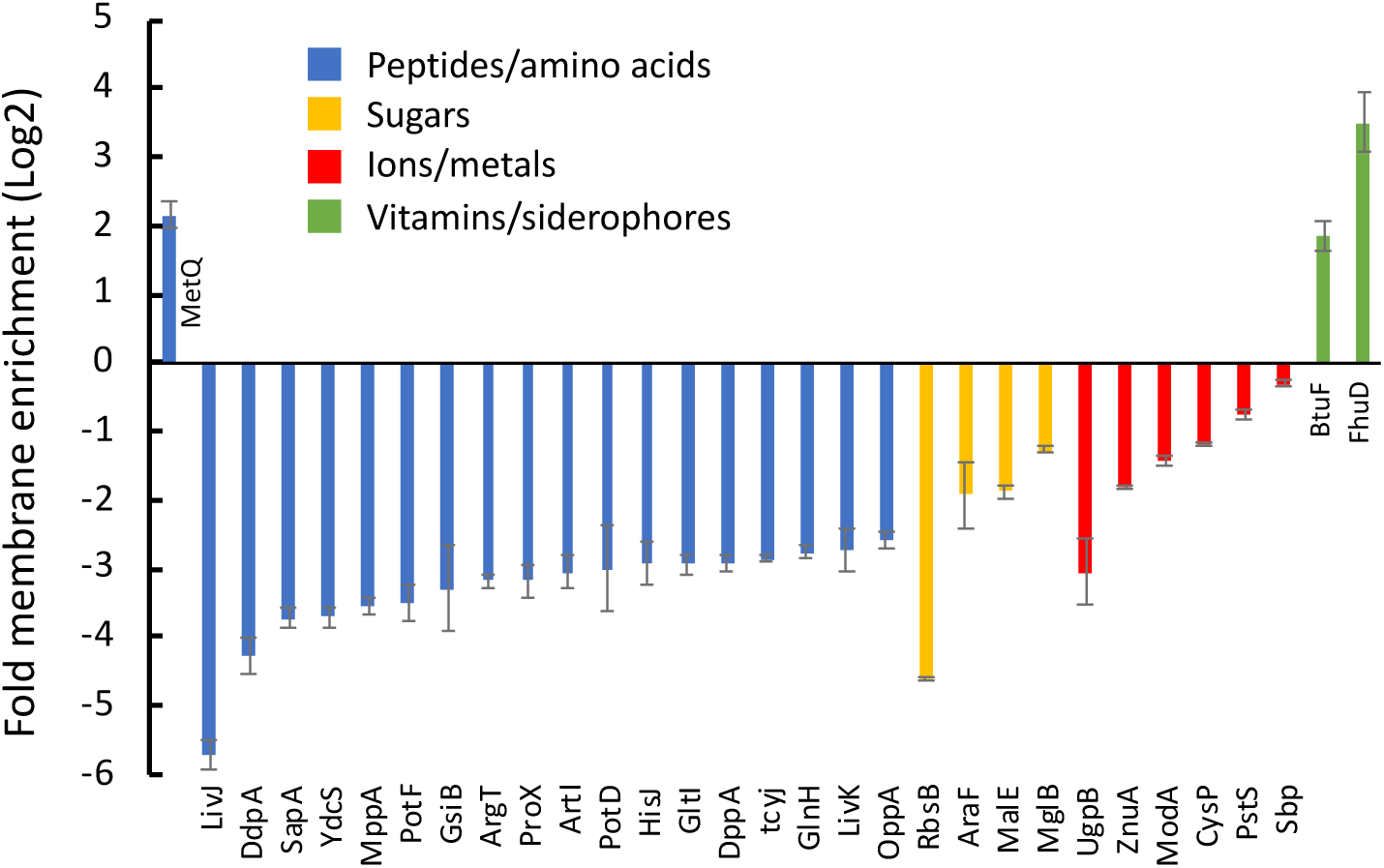
SBPs of Type-II systems are membrane bound. Shown (in Log2 scale) are the ratios between the abundances of the SBPs in whole cell lysates vs. their abundances in the membrane fraction. Results are means of biological triplicates and error bars represent standard deviations.

### Interdependence of expression and localization

The membrane association of BtuF and FhuD could be a result of a direct interaction with membrane lipids (as is the case of MetQ), a specific interaction with their cognate TMDs, or interactions with an unknown membrane protein(s). To distinguish between these possibilities, we repeated the experiments with strains carrying chromosomal deletions of the cognate TMDs of 8 different import systems. In terms of total expression levels, the SBPs were not deleteriously affected by the deletion of their cognate TMDs (Figure 6A). Notably, the expression of the SBPs for molybdate (ModA) and zinc (ZnuA) increased 20 and 30-fold (respectively) upon deletion of the corresponding TMDs. We suspect that this represents a substrate-specific starvation response. The cellular requirement both zinc and molybdate cannot be compensated for by alternative systems. In contrast, SBPs that were not induced by deletion of their TMDs are either of systems that can be complemented by alternative sugars/amino acids/peptides (*e.g.,* His, Mal, Yec, Met) or are not essential under these growth conditions (Fhu, Btu).

**Figure 6.**
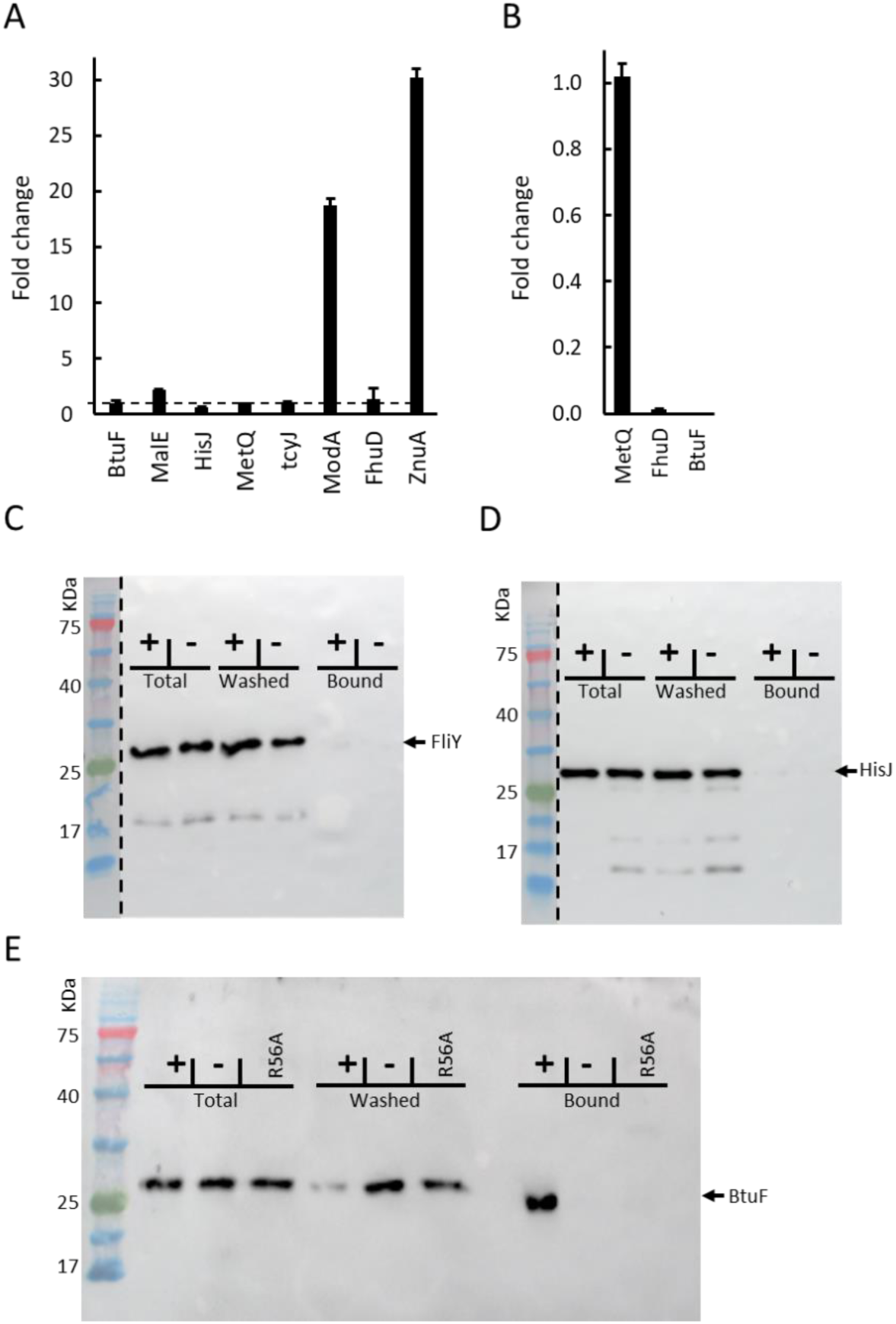
Co-localization of SBPs and TMDs in Type-II Systems. (A-B) Shown is the fold change in the abundances of the indicated SBPs following deletion of their cognate TMDs measured in total cell lysates (A) and membrane fractions (B). Shown are fold changes Error bars represent standard deviations, and results represent the mean of biological triplicates. (C-E): Cells deleted of the indicated TMDs were transformed with an empty plasmid or one harboring the deleted TMD (-, **+**, as indicated). Cells were then permeabilized by mild osmotic shock and purified SBPs were added at concentrations x10 their *K_D_* of interaction with their cognate transporters. Following a 15-min incubation, bound and unbound SBPs were separated by a wash step. Shown are immunoblots of SDS-PAGE of the total, washed, and cell-bound fractions for the Type-I systems Yec (C) and His (D) and for the Type-II system Btu (E). For the Btu system, an additional negative control was provided by cells that expressed the R56A mutant of BtuC which does not interact with BtuF (80).

We next tested the effects of the TMD-deletions on the membrane localization of the membrane associated SBPs. As shown in Figure 6B, the membrane localization of the lipid-anchored SBP MetQ was unaffected by deletion of its cognate TMD, MetI, confirming that its membrane association does not depend on its interaction with its cognate transporter (66). In contrast, the membrane association of BtuF and FhuD (Type-II SBPs) was completely abolished by deletion of their cognate TMDs (Figure 6B), demonstrating that their membrane association fully depends on specific interactions with their cognate transporters. To complement these studies, we performed membrane association biochemical assays. In these experiments we permeabilized the outer membrane of the cells by a mild osmotic shock treatment (see methods). We then added purified SBPs, incubated, and washed the cells. Our rational was that SBPs that freely diffuse in the periplasm will be removed by this wash step while those that are firmly membrane-associated will not. We tested two Type-I SBPs HisJ and FliY (43,68) and observed that they were indeed removed by the wash step (Figure 6C-D), in line with the proteomics data that shows that SBPs of Type-I systems are not membrane associated. In contrast, and in line with the membrane enrichment observed for Type-II SBPs (Figure 4), BtuF remained membrane-associated despite the wash step (Figure 6E). Also in line with the proteomics data was the finding that the membrane association of BtuF fully depended on the expression of its cognate TMDs (Figure 6E).

Like the substrate-binding proteins (SBPs), the nucleotide-binding domains (NBDs) did not exhibit detrimental effects on total expression levels upon the deletion of their corresponding transmembrane domains (TMDs). Contrarily, the majority of NBDs demonstrated an increase in expression when their TMDs were absent (Figure 7A). The only exception was observed with MetN, which completely failed to express in the absence of MetI. This, coupled with the distinct membrane tethering of MetQ, indicates that the methionine system is an exceptional case. Despite the heightened expression levels, the NBDs could hardly be detected in the membrane fraction (Figure 7B). This implies that, while the NBDs do not necessitate their corresponding transmembrane domains (TMDs) for expression per se, they are absolutely dependent on them for membrane association.

**Figure 7.**
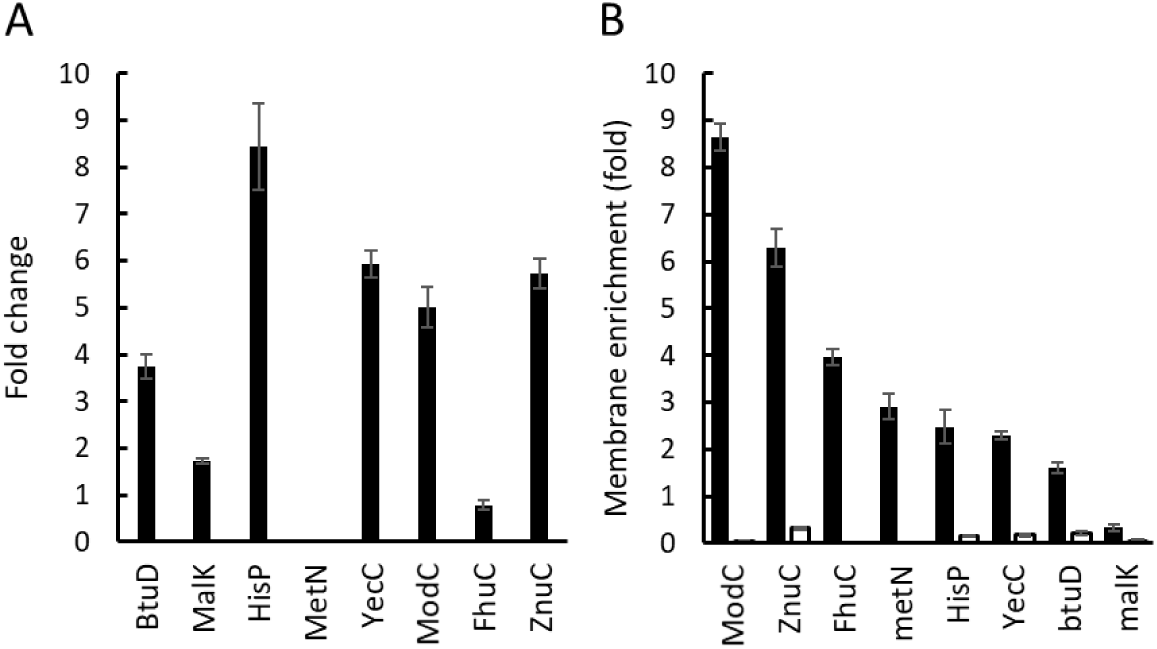
TMD-dependent localization of the NBDs. Shown is the fold change in the abundances of the indicated NBDs following deletion of their cognate TMDs, as measured in total cell lysates (A). Shown is the membrane enrichment fold of the indicated NBDs measured in WT or DTMD cells (full and open bars, respectively). Error bars represent standard deviations, and results represent the mean of biological triplicates.

Our collection of deletion strains included two transporters with heterodimeric TMDs: the histidine transporter HisP_2_QM and the maltose transporter MalFGK_2_. The structures of MalFGK_2_ (69–71) show that MalF and MalG are highly asymmetric, each contributing a different subset of residues to the translocation pore and to the interaction surface with the SBP. The consensus is therefore that this heterodimeric organization is an inherent and essential property of such transporters. To our great surprise, MalF was still present in the membranes of the Δ*malG* strain, and as if to compensate for the *malG* deletion, the amount of MalF increased almost exactly 2-fold (Figure S5A). In contrast, we could not detect MalG in the membranes of cells deleted of *malF* (not shown). These results suggest that MalF can insert into the membrane independently of MalG, and not *vice versa*. In line with these observations, the membrane association of MalK (the NBD of the maltose transporter) was hardly affected by the absence of MalG, but was nearly completely abolished in the absence of MalF, indicating that the expression of MalF is sufficient and essential for the membrane association/localization of MalK (Figure S5B). A somewhat similar phenomenon was observed with the histidine transporter HisPQM. Both HisM and HisQ (the TMDs) were detected in the absence of their counterpart. Of the two, HisQ was less dependent on HisM than *vice versa*, and its expression in the membranes of *ΔhisM* cells was ∼5-fold higher than that of HisM in *ΔhisQ* cells (Figure S5C). Strikingly, HisP (the NBD) was readily detected in membranes devoid of HisM but was completely absent from those devoid of HisQ (Figure S5C). The above observations the raise the possibility that these heteromeric transporters may also form stable homomeric membrane-bound complexes (*i.e.,* MalF_2_K_2_ and HisQ_2_P_2_), a suggestion that requires further investigation.

### Mechanistic implications of SBP-transporter stoichiometries

As shown in Figure 4, all Type-I systems exhibited a stoichiometric excess of the substrate-binding protein (SBP) in the range of 10-200-fold over the transporter. Notably, in the case of the Type-I maltose transporter, this surplus of the SBP has been established as crucial for activity. Mutations in the signal sequence of the SBP (MalE), which reduce its export to the periplasm and consequently lower its periplasmic copy number, lead to a decrease in transport rates. This reduction in transport rates results in impaired utilization of maltose as a carbon source (72,73). It appears, therefore, that in systems involved in the transport of abundantly available biomolecules, particularly those required in substantial quantities such as carbohydrates and amino acids, the stoichiometric excess of the SBP represents an essential mechanistic feature.

Surprisingly, and perhaps counterintuitively, Type-II systems exhibited a reversed ratio, with the transporter found in 10-100-fold stoichiometric excess over the substrate-binding protein (Figure 4). We postulated that this ratio might be substrate-modulated, shifting to the anticipated excess of SBP in the presence of substrate. However, our investigation revealed that the expression levels of BtuC, BtuD, and BtuF, as well as the stoichiometric ratio between them, remained unaffected by the presence of vitamin B12 (Figure S6). In contrast, acting as an internal control and consistent with prior findings (74), the expression of BtuB (the vitamin B12 outer membrane receptor/transporter) dropped to undetectable levels in the presence of vitamin B12 (Figure S6).

To assess the functional significance of the “reversed stoichiometry” observed in Type-II systems, we manipulated the transporter/SBP ratio within the Btu system by elevating the expression of the substrate-binding protein (SBP) (see methods for details). Through LC-MS^2^ analysis, we confirmed that the manipulated condition, designated as BtuF_high_, indeed replicates the substrate-binding protein (SBP)-transporter stoichiometric ratio observed in Type-I systems, corresponding to an approximately 100-fold excess of SBP.

In media where vitamin B12 uptake is non-essential for growth, we observed no distinction between wild-type (WT) cells expressing the intrinsic low levels of BtuF (referred to as BtuF_low_) and those expressing BtuF_high_ (Figure 8A). Subsequently, we replicated these experiments under conditions where methionine synthesis relies entirely on vitamin B12 uptake, rendering the transport activity of BtuCD-F crucial for growth (61). As shown in Figure 8B, in the presence of excessively high concentrations of vitamin B12 supporting 100% growth (5 nM), the elevated expression of the substrate-binding protein (SBP) did not confer either a growth advantage or a disadvantage.

**Figure 8.**
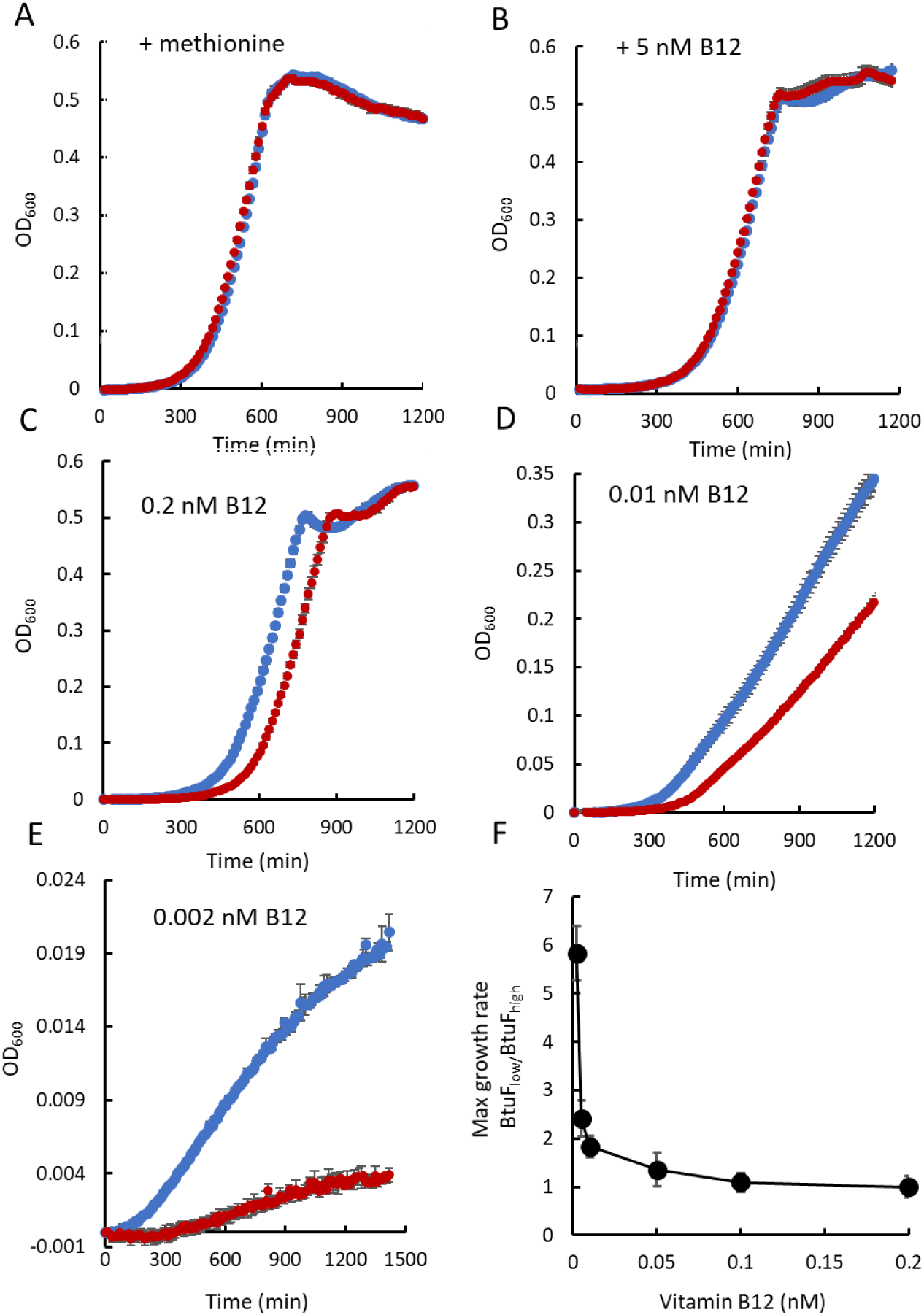
Unexpected advantage of SBP-Transporter stoichiometry < 1. *ΔmetE* cells rely on vitamin B12-dependent enzymatic methionine synthesis. Cell growth is contingent on exogenous methionine or vitamin B12, with BtuCD-F facilitating the import of the latter (27, 87). *ΔmetE* cells were cultured with the indicated additives, and the growth of cells with natural (BtuF_low_, blue curves) and elevated (BtuF_high_, red curves) BtuF levels is shown. In panel F, the ratio of the maximal growth rate between BtuF_low_ and BtuF_high_ cells is presented as a function of vitamin B12 concentrations. Notably, under vitamin B12 limitation, BtuF_low_ cells exhibit approximately six-fold faster growth than BtuF_high_ cells; under vitamin B12 abundance, growth is identical. Results are averages of biological triplicates, with error bars indicating standard deviations.

Remarkably, as we lowered the concentrations of vitamin B12, making it limiting for growth, the BtuF_low_ cells exhibited a significant growth advantage, and this advantage became more pronounced as the concentrations decreased (Figure 8C-F). This stands in stark contrast to the observations for the model Type-I transporter MalFGK-E, where reduced substrate-binding protein (SBP) led to decreased transport and impaired growth (72,73).

This unexpected phenomenon, where increasing amounts of one of the components of a system leads to reduced output, can be explained by considering the physiologically relevant concentrations of the system’s components and their absolute numbers. As established in previous observations (61,75), and further corroborated by our findings (Figure 8B), as little as 5 nM vitamin B12 supports maximal growth.

This concentration corresponds to a periplasmic presence of approximately 3-4 vitamin B12 molecules, perfectly aligning with the cellular copy number of BtuF (4 copies/cell, Table S3). The cellular copy number of the transporter is approximately 20 (Table S3), indicating that transporter docking sites are available for all vitamin B12-bound BtuF molecules (Figure 9A). In the manipulated system (BtuF_high_), the expression of excess BtuF relative to vitamin B12 leads to the generation of a population of vitamin B12-free BtuF molecules. Previous studies have demonstrated that vitamin B12-free BtuF exhibits very high affinity towards its cognate transporter (47). Consequently, these ligand-free BtuF molecules will compete with the fewer ligand-bound ones for docking to the available transporters (Figure 9B). Given their extremely slow dissociation rate (∼10^−7^ S^−1^, (47)) this will effectively clog the system and reduce uptake rates, as shown in Figure 8C-F and schematically depicted in Figure 9B. A similar problem does not arise in Type-I systems. Their substrates are sufficiently abundant to saturate the SBPs, diminishing the competition between substrate-free and loaded SBPs (Figure 9C-D). In addition, Type-I transporters preferentially associate with substrate-loaded SBPs, and their SBPs rapidly dissociate from the transporters, avoiding the problem of clogging the transporters with substrate-free SBPs (44–48). It therefore seems that two different SBP: transporter stoichiometries evolved, with ratios >>1 and <<1 tailored for acquiring biomolecules of high and low abundance, respectively.

**Figure 9.**
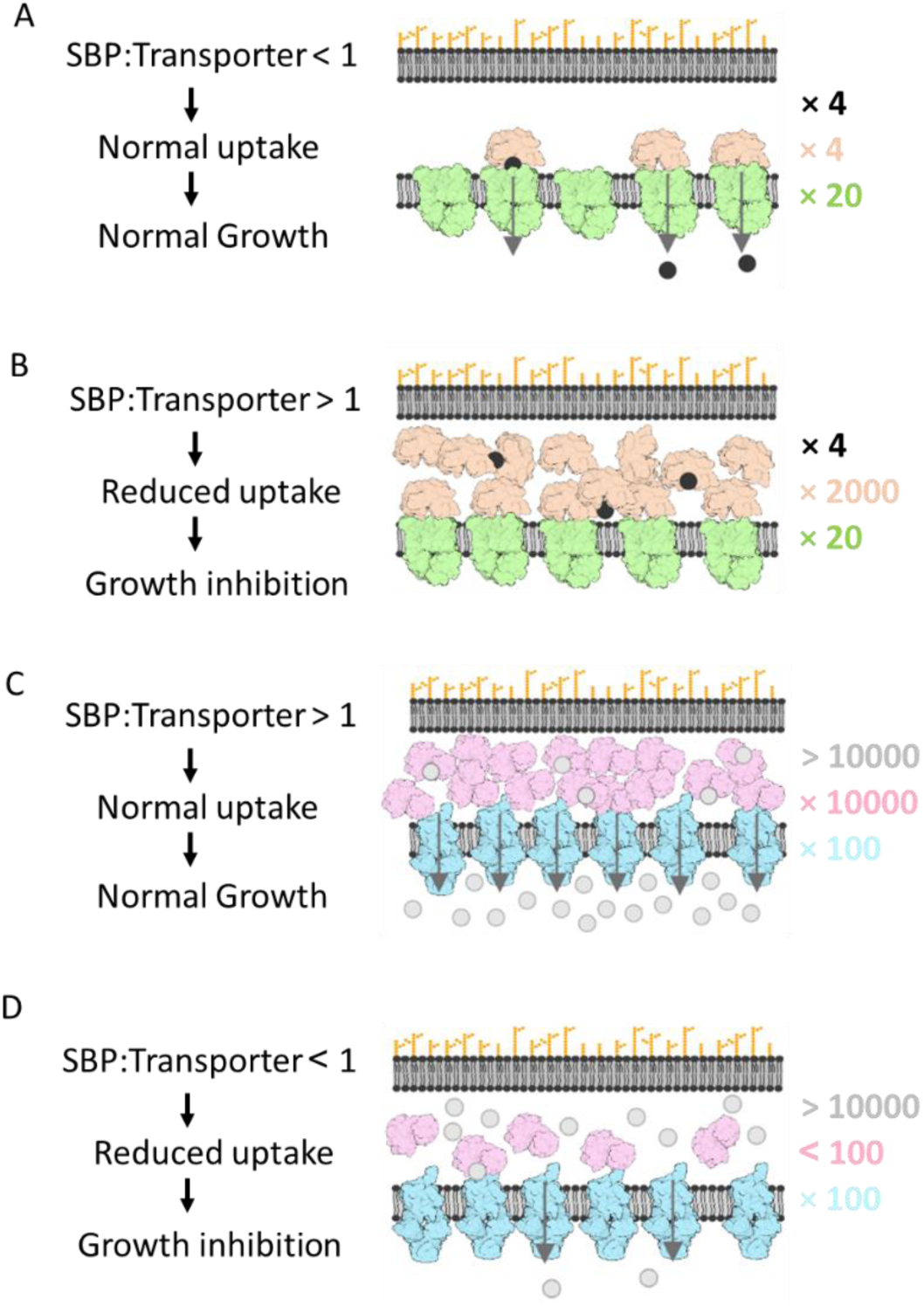
A model for mechanistic adaptation at the cellular level. Systems that import rare compounds such as vitamin operate optimally when the SBP is present at stoichiometric paucity relative to the transporter (A). Because of the low amounts of environmental ligand, the presence of excess SBP will lead to a substantial population of ligand-free SBPs which will compete with the fewer ligand bound ones in docking to the available transporter molecules (B). Conversely, systems that import abundant compounds, such as amino acids, operate optimally when the SBP is present at stoichiometric excess relative to the transporter (C). In such systems, the excess of the SBP does not present a problem due to the inherent instability of the transporter-SBP complex and the preferential interaction of the transporter with the substrate-loaded SBP. However, in such systems, stoichiometric paucity of the SBP will lead to fewer ligand binding events, fewer fruitful transporter-SBP interaction events, less transport, and reduced fitness (D).

In summary, we integrated proteomic analysis with functional assays to explore correlations between thermodynamic/kinetic properties, substrate specificity, copy number, and component stoichiometry in ABC import systems. Our findings reveal abundances that vary over 4-5 orders of magnitude and correlate with the quantities of nutrients requiring acquisition. Additionally, we observed that receptors (SBPs) of systems importing rare compounds do not freely diffuse in the periplasm but are firmly docked to their cognate membrane-embedded transporter. Most intriguingly, such systems utilize a “reversed” stoichiometry where the receptor is found in paucity relative to the transporter. Taken together, the results showcase correlations between physiological roles, molecular-level attributes, and cellular-context parameters, suggesting their coevolution to optimize the output of multi-component systems.

## Materials & Methods

### Mass Spectrometry sample preparation and data analysis

Cultures of the *E. coli* K12 strain BW25113, the progenitor strain of the Keio clone collection (32), were grown to mid-exponential phase in M9-glucose medium, harvested by centrifugation, washed in PBS and stored in −80. Where indicated, membranal, cytosolic, and periplasmic fractions were prepared from these cell pellets. For preparation of membrane fraction, cells were re-suspended in 50 mM Tris-HCl, pH 7.5, 0.5 M NaCl, and ruptured by tip sonication (3 × 20 s, 600 Watts). Debris and unbroken cells were removed by centrifugation (10 min, 10,000 × *g*) and the membranes were pelleted by ultracentrifugation at 120,000 × *g* for 45 min. The membranes were washed and re-suspended in 50 mM Tris-HCl, pH 7.5, 0.5 M NaCl and stored in −80 °C. Where indicated, membranes were subjected to an additional wash step with 100mM sodium bicarbonate, 2 M sodium chloride, or 25 mM EDTA.

Next, the samples were dissolved in 10mM DTT 100mM Tris and 5% SDS, or 1 % DDM, 1% DM or a mixture of or 1% DDM + 1% DM, as indicated. Samples were sonicated and boiled in 95^0^ C for 5 min and precipitated in 80% acetone. The protein pellets were dissolved in in 9M Urea and 400mM ammonium bicarbonate than reduced with 3mM DTT (60°C for 30 min), modified with 10mM iodoacetamide in 100mM ammonium bicarbonate (room temperature 30 min in the dark) and digested overnight in 2M Urea, 25mM ammonium bicarbonate at 37°C using a 1:50 (mol/mol) of modified trypsin (Promega), 1:10 (mol/mol) Chymotrypsin)Roche), 1:50 (mol/mol) ProAlanase (Promega), 1:50 (mol/mol) WalP (Cell signaling technology), 1:3 (mol/mol) Pepsin(Promega), or 1:50 (mol/mol) Lys-C(Fujifilm), as indicated.

The peptides were desalted using C18 tips (Top tip, Glygen), dried and re-suspended in 0.1% Formic acid, and resolved by reverse-phase chromatography on 0.075 X 180-mm fused silica capillaries (J&W) packed with Reprosil reversed phase material (Dr Maisch GmbH, Germany). The peptides were eluted with linear gradients of acetonitrile with 0.1% formic acid in water: 120 minutes of 5 to 28%, 15 minutes of 28 to 95%, and 25 minutes at 95% at flow rates of 0.15 μl/min. Mass spectrometry was performed using a Q Exactive HFX mass spectrometer (Thermo) in a positive mode (m/z 300–1800, resolution 120,000 for MS1 and 15,000 for MS2) using repetitively full MS scan followed by collision induces dissociation (HCD, at 27 normalized collision energy) of the 18 most dominant ions (>1 charges) selected from the first MS scan. The AGC settings were 3×106 for the full MS and 1×105 for the MS/MS scans.

The intensity threshold for triggering MS/MS analysis was 1×104. A dynamic exclusion list was enabled with exclusion duration of 20 s.

Mass spectrometry data was analyzed using the MaxQuant software 1.5.2.8 (76) for peak picking and identification using the Andromeda search engine, searching against the human proteome from the Uniprot database with mass tolerance of 6 ppm for the precursor masses and 20 ppm for the fragment ions. Oxidation on methionine and protein N-terminus acetylation were accepted as variable modifications and carbamidomethyl on cysteine was accepted as static modifications. Minimal peptide length was set to six amino acids and a maximum of two miscleavages was allowed. The data was quantified by label free analysis using the same software. Peptide- and protein-level false discovery rates (FDRs) were filtered to 1% using the target-decoy strategy. Protein table were filtered to eliminate the identifications from the reverse database, and common contaminants and single peptide identifications Statistical analysis of the identification and quantization results was done using Perseus 1.6.10.43 software (77).

### Calculation of cellular copy numbers

To calculate cellular copy numbers, equation 1 was used to determine relative iBAQ values (40,41) for each protein:

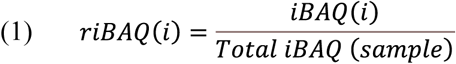

where *riBAQ(i) is the* relative iBAQ value of protein *i*, *iBAQ(i)* is its iBAQ value as determined by MaxQuant, and *Total iBAQ (sample)* is the sum of all iBAQ values of the sample.

Using equation 2, the mass (in grams) was calculated for each protein by multiplying *riBAQ(i)* by the total protein mass of the sample:

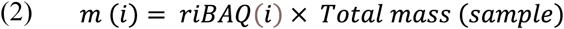

The total number of moles present in the sample for a given protein was obtained using equation 3,

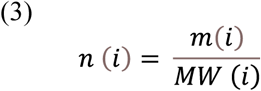

where *n* is the total number of moles, *m* is the total mass of a given protein present in the sample, and *MW* is its molecular weight. The total number of molecules of a given protein present in the sample, *N (i)*, was then obtained using Avogadro’s number (equation 4).

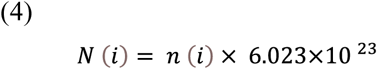

Finally, cellular copy numbers, *C(i)*, were determined using equation 5

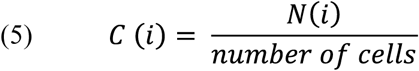

Where *N(i)* is the total number of molecules obtained using equation 4 and the number of cells determined by colony counting.

### Membrane association assays

For the membrane association studies only fresh cultures, ones that have not been frozen and/or stored, were used. The experiments were conducted essentially as previously described (47,78) with the following modifications. Cells were transformed with an empty vector or with the same vector encoding the transporters YecSC, HisPQM, or BtuCD, as indicated. Cultures were grown in LB-ampicillin medium at 37 °C with shaking to mid-exponential phase, and protein expression was induced by isopropyl β-D-1-thiogalactopyranoside (0.5 mM) for 1 hour. Cells were harvested and resuspended in 10 mM Tris⋅HCl, pH 7.5, and 0.75 M sucrose. The cell suspension was then incubated on ice for 20 min in the presence of 100 µg/mL lysozyme (Amresco) and 2 volumes of 1.5 mM EDTA. The spheroplasts were then stabilized by addition of 25 mM MgCl_2_ and DNA originating form ruptured cells was digested with 100 µg/mL DNase-I (Worthington). The spheroplasts were pelleted and resuspended in 100 mM Tris⋅HCl, pH 7.5, 150 mM NaCl, and 5 mM MgCl_2_ to an OD_600_ of about 10 and kept on ice until use. Purified FLAG-tagged SBPs were added to the spheroplasts suspension (160 nM BtuF, 5mM FliY, 15mM HisJ) and following a 10-min incubation, unbound material was removed by 5 min centrifugation at 2500 ×g. The amount of FLAG-tagged SBPs in the total, washed, and bound fractions was visualized using standard immuno-blot procedures, using an anti-FLAG M2 HRP-conjugate antibody (Sigma).

### Vitamin B12 utilization assays

This assay is based on the original protocol developed by the Kadnar lab (79). In the *E. coli ΔmetE* strain methionine biosynthesis is solely catalyzed by MetH, a vitamin B12-dependent methionine synthase. In this strain growth is therefore dependent on the exogenous addition of either methionine or vitamin B12. When the latter is supplied, growth fully depends on the uptake function of BtuCD-F (80). In the experiments shown in Figure 5, BtuF_low_ cells refer to *ΔmetE* cells transformed with an empty vector, and BtuF_high_ cells refer to the same cells transformed with a BtuF-encoding vector. Expression of the plasmid was not induced during the experiment, and hence, excess BtuF in the BtuF_high_ cells originates from promoter leakiness. As detailed in the text, this leaky expression amounts to 1000-2000 copies of BtuF per cell. Cultures were grown in LB media supplemented with 50 μg/mL kanamycin and 100 μg/mL ampicillin to mid log phase, washed and resuspended in Davis minimal media to OD_600_ of 0.05. Cultures (0.2 mL) were then grown in absence or presence of methionine or vitamin B12, as indicated. The optical density of the cultures was measured every 5 min for 12 h using an automated plate reader (Infinite M200 Pro; Tecan).

## Acknowledgements

This work was supported by grants from NATO Science for Peace and Security Program (SPS project G5685), and the Israeli Academy of Sciences project 1006/18.

All data needed to evaluate the conclusions in the paper are present in the paper and/or the Supplementary Materials.

## Supplementary legends

**Supplementary Table S1.** Listed are the systems suggested to function as ABC import systems (1). The three systems that were excluded from the analysis, and the reasons for their exclusion, are highlighted in yellow.

**Supplementary Table S2.** Shown are the different sample preparation protocols used, indicating strain (WT or deletion mutants), cellular fraction (where applicable), detergent, and protease. The components of the ‘ABC importome’ identified in each experiment are indicated by **√** or **-**

**Supplementary Table S3.** The complied dataset of the components of the ‘ABC importome’ identified in this study. Results are means of at least three biological replicates.

**Supplementary Figure S1.**
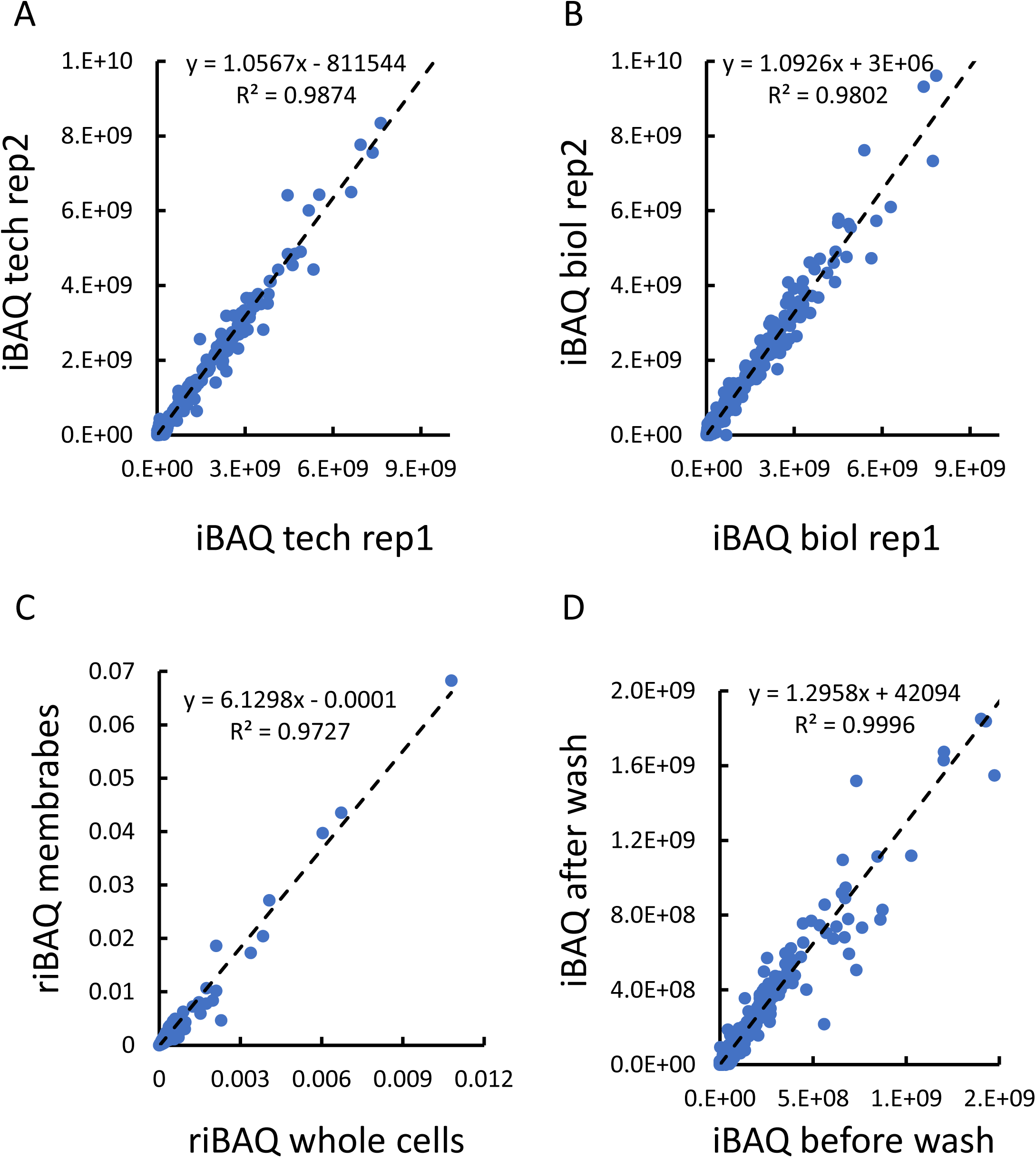
Examples of high correlation between different datasets. Shown are the correlations between the iBAQ values obtained for two technical replicates (A), two technical biological replicates (B), for whole cells and membranes fractions (C), and for membranes before or after washing with 100 mM sodium carbonate (D). Also shown are the linear regressions (dashed black lines), their slopes, and the Pearson correlation coefficients.

**Supplementary Figure S2.**
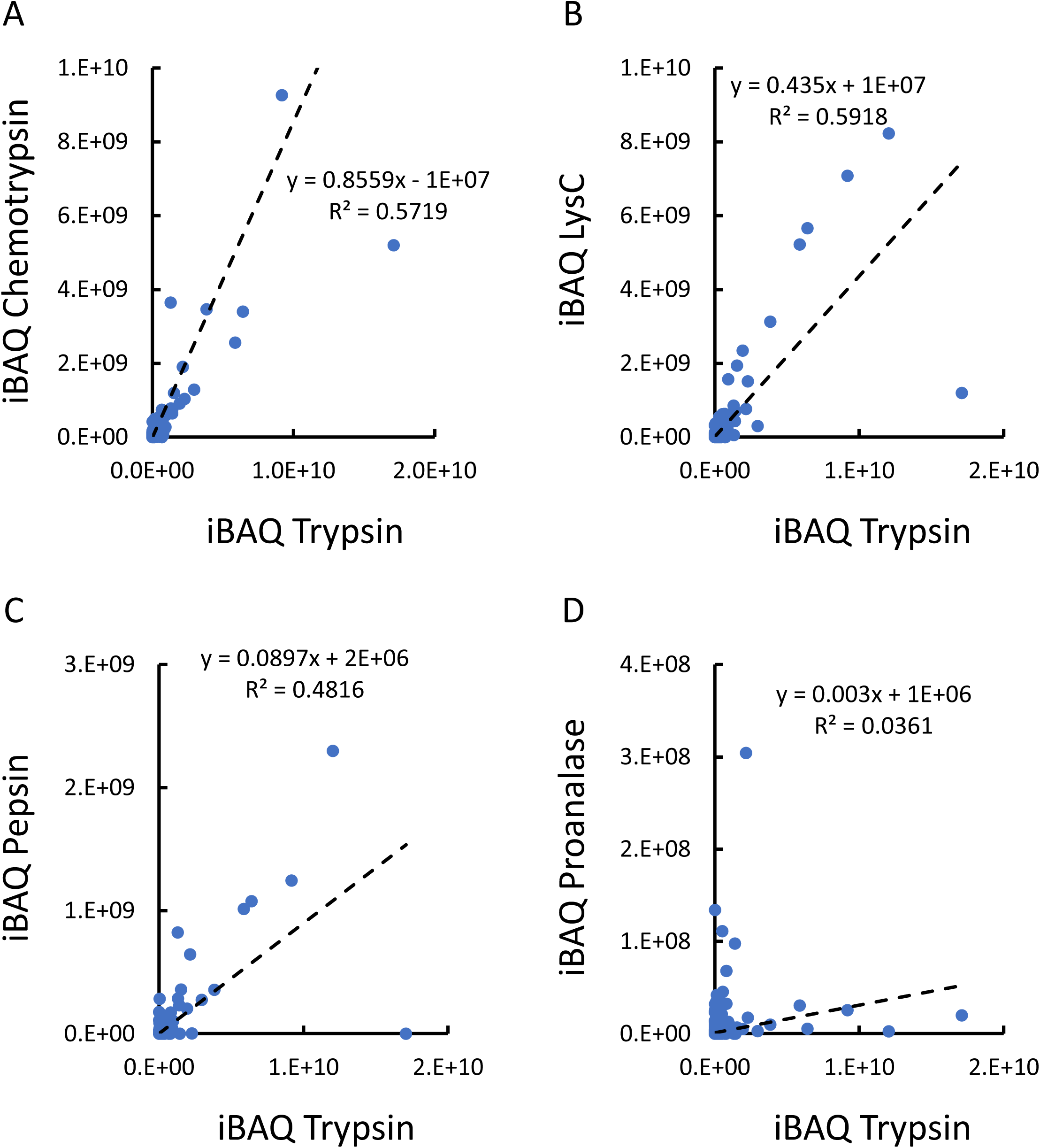
Low correlation between samples digested with different proteases. Shown are the correlations between the iBAQ values obtained for samples digested with either trypsin or chemotrypsin (A), digested with either trypsin or LysC (B), digested with either trypsin or pepsin (C), and digested with either trypsin or Proanalase (D). Also shown are the linear regressions (dashed black lines), their slopes, and the Pearson correlation coefficients.

**Supplementary Figure S3.**
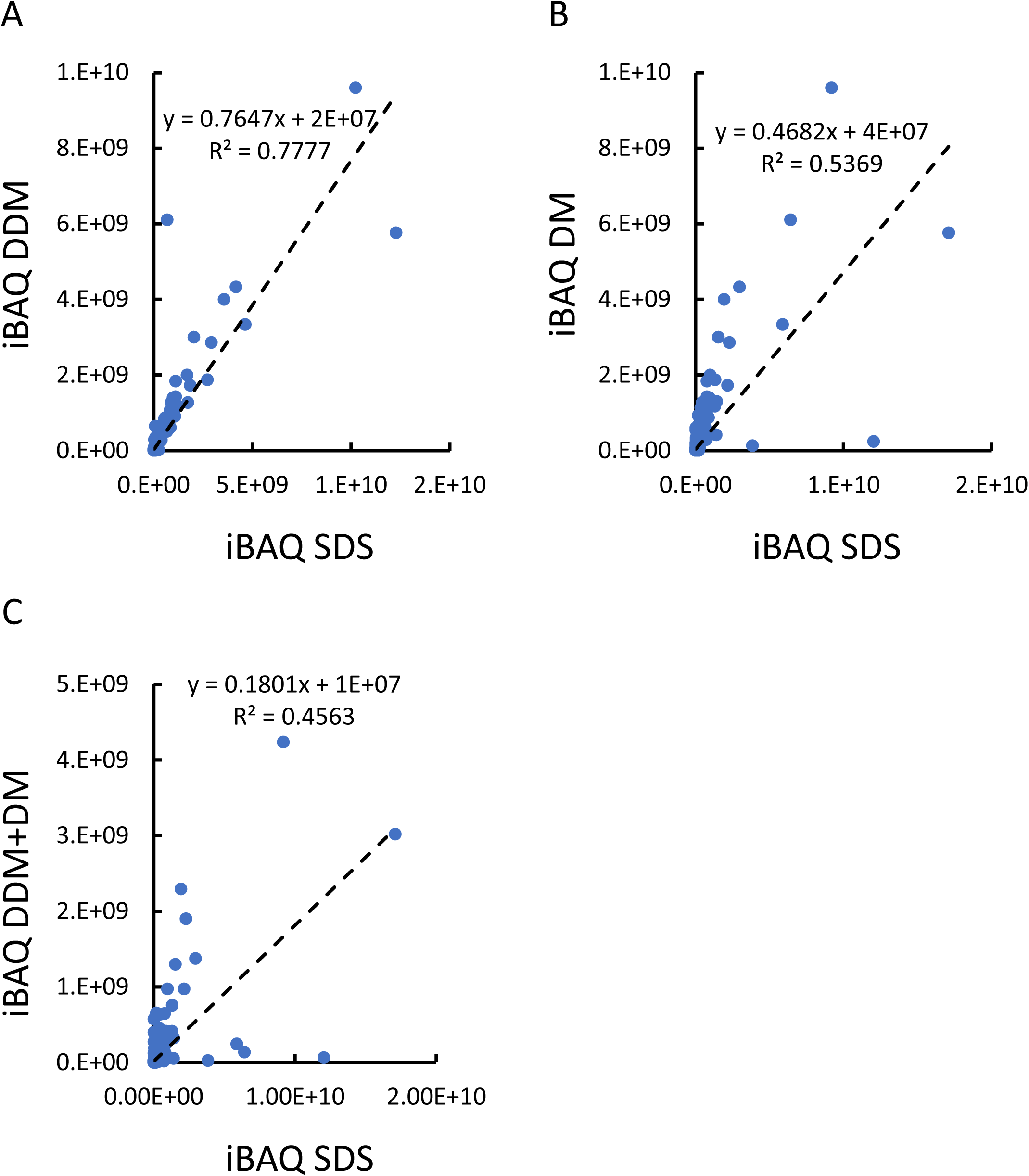
Low correlations between samples extracted with different detergents. Shown are the correlations between the iBAQ values obtained for samples extracted with either sodium dodecyl sulfate (SDS) or n-dodecyl β-D-maltoside (DDM) (A), SDS or n-decyl β-D-maltoside DM (B), and SDS or a mixture of DDM+DM (C). Also shown are the linear regressions (dashed black lines), their slopes, and the Pearson correlation coefficients.

**Supplementary Figure S4.**
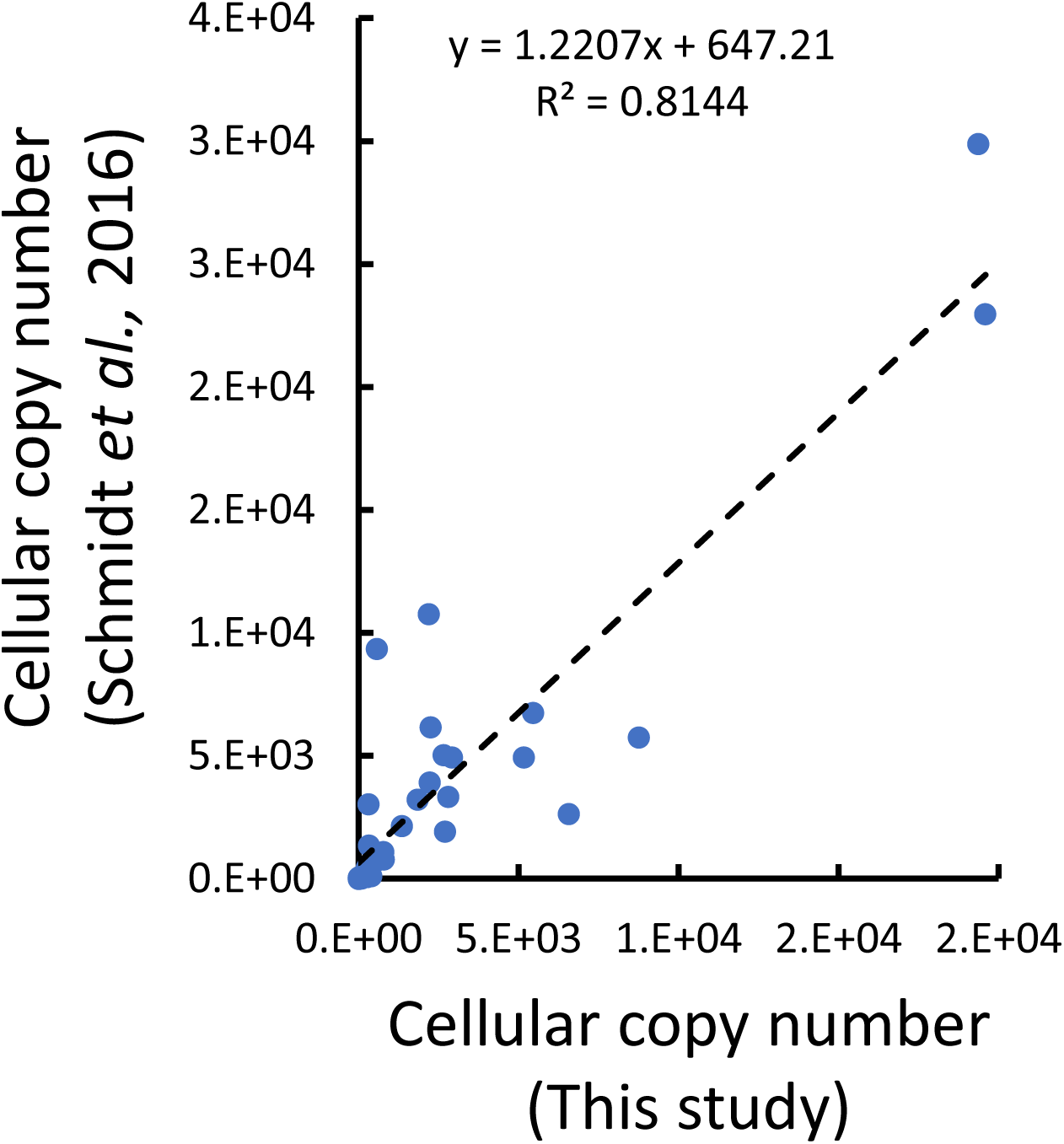
A comparison between the cellular copy numbers of SBPs determined by Schmidt et al (2) and in this study. Also shown is the linear regression (dashed black line), its slope, and the Pearson correlation coefficient.

**Supplementary Figure S5.**
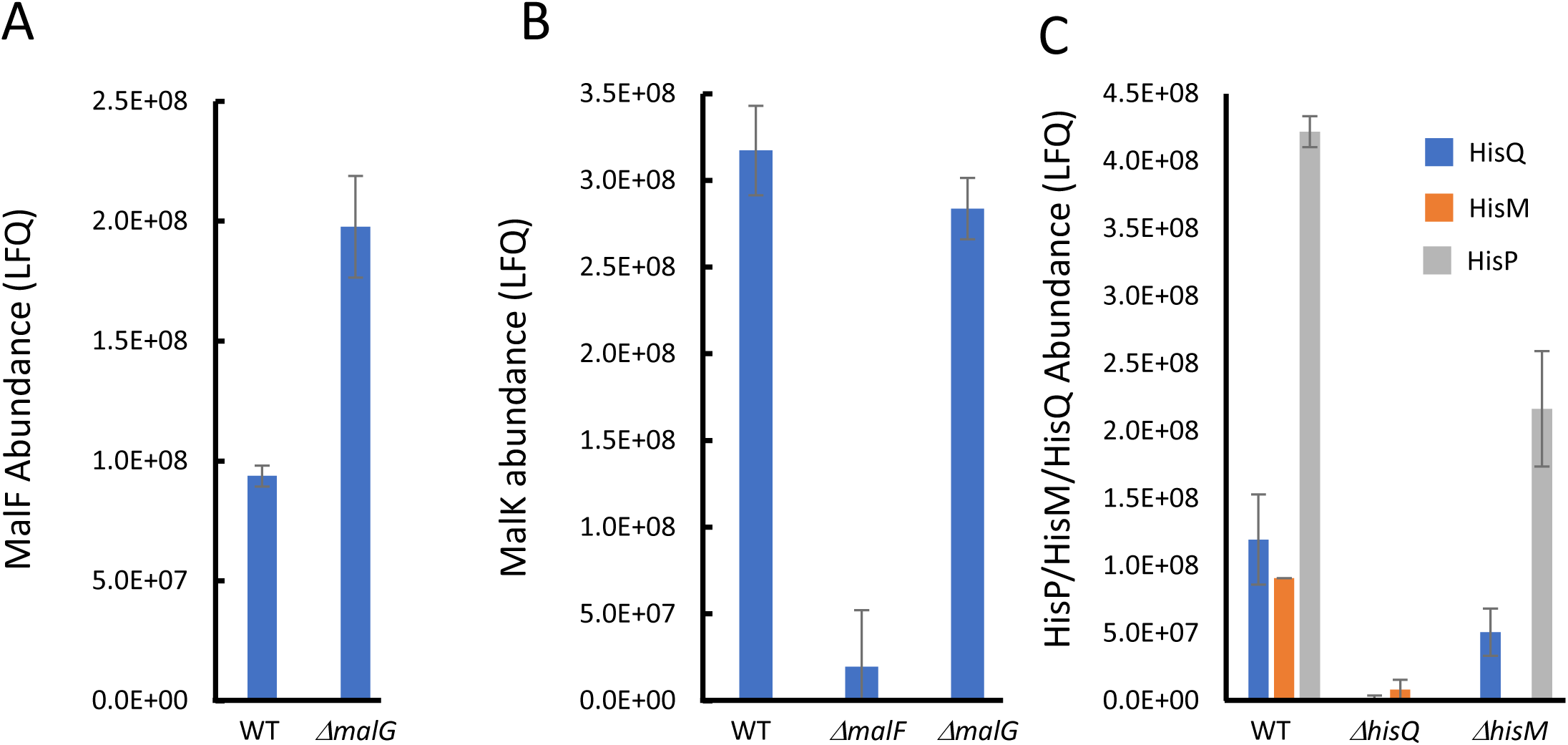
Asymmetric biogenesis of heteromeric systems. Shown are the membrane fraction abundances of MalF in WT and *ΔmalG* cells (A), of MalK in WT, *ΔmalF*, and *ΔmalG* cells (B), and of HisP, HisM, and HisP (as color coded) in WT, *ΔhisQ*, and *ΔhisM* cells (C). Results are means of biological triplicates and error bars represent standard deviations.

**Supplementary Figure S6.**
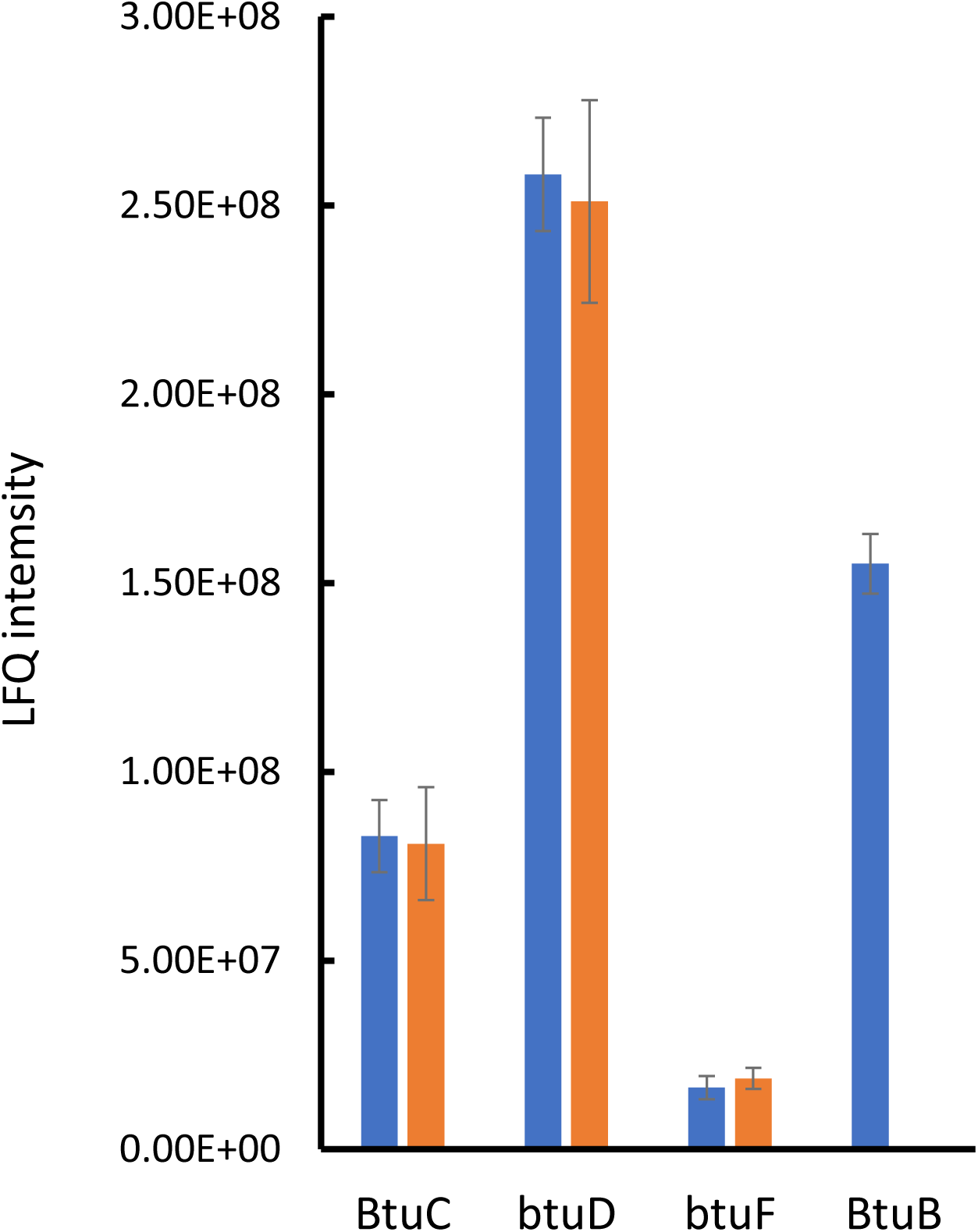
The transporter-SBP stoichiometry of BtuCD-F in unaffected by vitamin B12. Shown are abundances of BtuC, BtuD, BtuF, and BtuB (as indicated) measured in cells grown in the absence or presence of 50 nM vitamin B12 (blue and orange bars, respectively). Results are means of biological triplicates and error bars represent standard deviations.

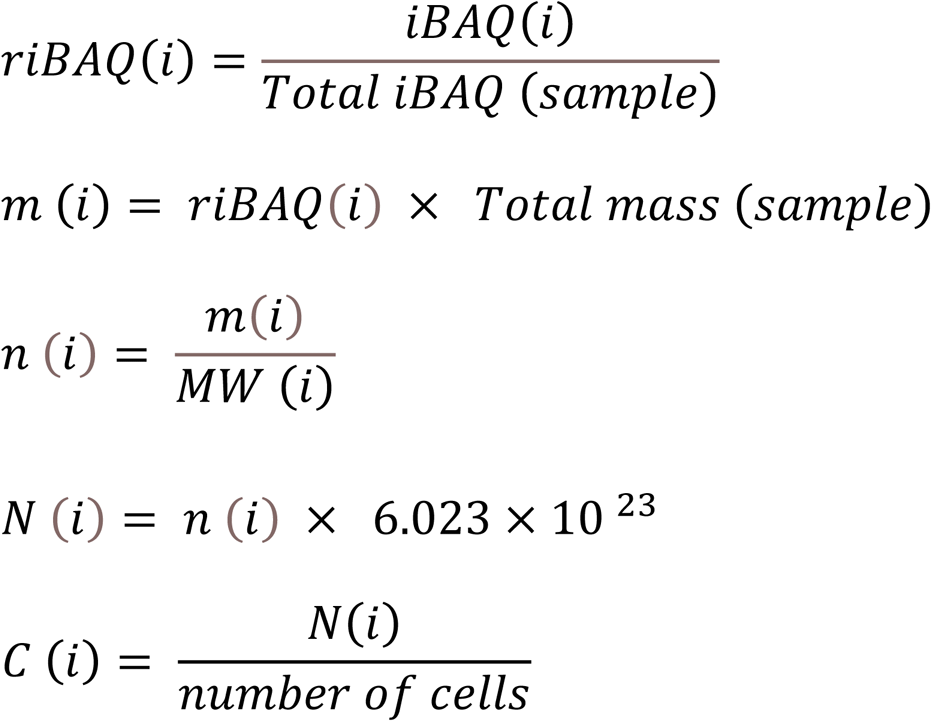

